# Functional disruption of *vgsc* reveals haplosufficiency with implications for insecticide resistance and genetic control in *Anopheles gambiae*

**DOI:** 10.64898/2026.09.09.750417

**Authors:** Poppy Pescod, Ashley Hall, Sian Wakley, Matt Craske, Louise Marston, Amalia Anthousi, Carla Siniscalchi, Josie Shepherd, Molly Kelly, Saffron Mackey, Kurtis Blinkhorn, Tony Nolan

## Abstract

Pyrethroid insecticides on bednets have been the mainstay of malaria control since the millennium by killing mosquitoes. The target of these pyrethroids is the voltage-gated sodium channel (VGSC). Resistance mutations at the pyrethroid binding site (L995F/S) have spread widely, but functional tools to dissect *vgsc* itself have lagged behind the population-genetic surveillance data. Manipulation of *vgsc* via knockout or functional mutation would be a vital step towards understanding its contribution to insecticide resistance, as well as exploring its utility as a genetic control target. Here we generate and characterise two CRISPR-edited *An. gambiae* lines: an exonic knockout (*vgscKO*) and an intronic CRISPR-mediated cassette exchange intermediate (*vgscInt*) sited near SNPs linked to L995F. Both insertions are viable and fertile in heterozygosity but homozygous lethal, demonstrating that *vgsc* is haplosufficient and therefore the locus represents a viable target for population suppression genetic control strategies. These insertions can also be used as balancers for studying *vgsc* mutations without the confounding effects of wild type alleles. We investigate the insecticide resistance phenotype of *vgscKO* and demonstrate that knocking out one copy of *vgsc* produces reduced susceptibility to deltamethrin and DDT at sub-discriminating doses, with implications for *vgsc*-mediated mechanisms of insecticide resistance.

**Author summary:** With malaria cases once more on the rise, current insecticide-based control strategies are being assessed and new interventions are being considered. Pyrethroid insecticides are the mainstay of vector control for malaria, but resistance at their target site the VGSC is reducing their effectivity. Genetic tools such as gene knockouts, including knock-ins which impair gene function, will allow investigation into the binding dynamics of pyrethroids to VGSC; but such tools in *Anopheles* still need developing. Here we present and characterise two new *Anopheles gambiae* lines with insertions at different loci within the *vgsc* gene. Both insertions produce knockouts of the *vgsc gene.* We find that *An. gambiae* is viable with one functional copy of *vgsc* but non-viable with no working copies, allowing use of either *vgsc* knockout as a balancer allele for investigating functional mutations of the gene. We also find that the *vgsc* knockout produces a low-level resistance to pyrethroid insecticides, suggesting pyrethroid resistance may be partially VGSC dose dependent. The *vgsc* gene merits exploration as a genetic control target as well as a mechanistic model system, due to its rare combination of high conservation, functional constraint, essentiality, and relevance to insecticide resistance.

## Introduction

Following the success and subsequent faltering of insecticide-treated bednets (ITNs) against the malaria vector *Anopheles*, malaria cases are once more rising across the globe (1). While new insecticides are under development, ITNs have been predominantly reliant on the pyrethroid class of insecticides due to their efficacy and safety profile (2, 3). Pyrethroids target the nervous system by binding to the voltage-gated sodium channel (VGSC) and preventing its transition from an active to an inactive state, depolarising the nerve membrane and paralysing the insect (4). The VGSC is made up of four homologous domains, each consisting of 6 transmembrane segments (S1-S6), which together form a transmembrane channel; the pyrethroid binding site sits within this channel (5, 6).

Whilst mammals have multiple *vgsc* genes encoding channels with different properties for specific cell types or developmental stages, insects rely on a single *vgsc* gene with alternative splicing and post-transcriptional RNA modification to achieve the same functional diversity (5, 7, 8). The *Anopheles gambiae vgsc* gene (AGAP004707) consists of 39 annotated exons (10-15 coding exons used in each final transcript) with 13 known splice variants, making it one of the most transcriptionally complex genes in the organism (9). Despite this structural versatility, strong purifying selection on this essential gene has resulted in a highly conserved amino acid sequence amongst insect species, and polymorphisms within the coding sequence of *Anopheles* are uncommon (9).

However, single nucleotide polymorphisms (SNPs) in the pyrethroid binding site can be enough to reduce binding and cause resistance to insecticides (10). In *Anopheles*, non-synonymous SNPs at the L995 amino acid codon within the pyrethroid binding site have produced two of the most potent and widespread target site resistance mutations to pyrethroids – L995F (leucine to phenylalanine) and L995S (leucine to serine). Secondary SNPs in the *vgsc* gene have also been identified, with some showing strong linkage disequilibrium with L995 mutations, implying an additive or compensatory effect on mosquito fitness (9). The examination of L995F/S haplotype backgrounds demonstrates that both mutations have arisen multiple times independently, with haplotype analysis demonstrating spread amongst species within the *An. gambiae* complex and penetration across geographical barriers (11).

The adversarial selection between *vgsc* sequence conservation and insecticide pressure makes developing genetic tools to understand the *vgsc* gene potentially difficult but important; it also makes *vgsc* an interesting potential target for genetic control strategies, such as gene drives. Gene drives are selfish genetic elements designed to insert themselves into specific sequences by cutting the target site and copying their own genetic material over, by design either reducing or altering a population at a rapid evolutionary timescale (12–14). To be effective gene drive target genes need to be highly conserved and, in the case of most suppression drives, haplosufficient – producing no fitness loss when knocked out in heterozygosity, but full loss of function when knocked out at homozygosity. The *vgsc* gene shows promise as a suppression target; but understanding the phenotypic and genetic impacts of altering the *vgsc* gene will be essential to understanding its potential for use in genetic control strategies as well as its role in insecticide resistance.

Gene knockouts, either through non-viable small mutations or knock-in of a cassette that disrupts coding sequence, can be used as balancers for accurately assessing the phenotypic impact of mutations in the alternate chromosome, and for checking whether a gene is haplosufficient. No knockouts of the *vgsc* gene are documented in *Anopheles*; in *Drosophila*, a knockout of the ortholog *para* produces a dominant impact on flight, seizure activity and behaviour, although no impact on insecticide resistance was tested (15).

As well as knockouts, functional single nucleotide modifications are an essential piece of the genetic toolkit to investigate gene function. This is particularly pertinent to the *vgsc* gene where SNPs are known to be capable of strong phenotypic effects and several SNPs of unknown function have been discovered tightly linked to the L995F mutation, indicating either an amelioration of fitness costs or an additive impact on insecticide resistance (9). Engineering single nucleotide edits without interfering with the function of the gene requires markerless editing – whereby the precise edit is achieved with no extra genetic material, such as a fluorescent marker, inserted into the gene. This is fully achievable via CRISPR-Cas9 or other technologies, but transformation rates are generally around 5% (16–18) and screening for successfully edited individuals is difficult and time consuming. CRISPR-mediated cassette exchange (CriMCE) is a technique involving insertion of a fluorescent placeholder as an intermediate step to making a markerless edit. The fluorescent placeholder is subsequently removed in a second injection round and replaced by homology-directed repair with the desired markerless edit that recapitulates the ‘field’ allele of choice, allowing screening for successful edits by the absence of the marker, which greatly increases screening efficiency (19). This approach is greatly facilitated when the placeholder can exist in homozygosity; as the *vgsc* gene is highly conserved and only one copy exists in insects, a homozygous knockout is unlikely to be viable. An intronic placeholder insert nearby SNPs of interest would in theory allow for viability of the modified *vgsc* gene as well as modification of nearby sequences via CriMCE.

In this work we have produced a non-functional allele of the *vgsc* gene in the malaria mosquito *An. gambiae* by knocking in a fluorescent protein into exon 19, and a CriMCE intermediate with a fluorescent protein situated in the intron between exons 30 and 31, nearby several SNPs in linkage disequilibrium with the L995F mutation. We have assessed the impact of *vgsc* knockout on fitness and fertility, as well as insecticide resistance status, and explored the viability of intronic CriMCE intermediates for *vgsc* SNP editing.

## Results

### Insertion and validation of fluorescent markers

For the *vgsc* knockout line (hereafter referred to as *vgscKO*), *An. gambiae* G3 strain embryos were microinjected with a helper plasmid containing a CRISPR-Cas9 cassette with a gRNA cutting on exon 19 (AGAP004707-RD), and a donor plasmid containing a cyan fluorescent protein (CFP) under Actin5c promotion, flanked by 0.9 kb homology arms (**Figure 1**). For the intronic CriMCE intermediate line with an intronic insertion (hereafter *vgscInt*), injections were performed in a previously injected *An. gambiae* line called *Kisumu-F/F*, derived from the Kisumu strain with a homozygous base change at the L995 site (TTA-> TTT) (17). *Kisumu-F/F* was microinjected with a CRISPR-Cas9 helper plasmid cutting on exon 31 (AGAP004707-RD), and a donor plasmid containing a dsRed protein under 3xP3 promotion (**Figure 1**). The homology arms on the donor plasmid (0.8kb) were designed to completely restore function to the cut exon, inserting the fluorescence cassette in the 5’ intronic region, and replacing the ‘CGG’ PAM site present in the right homology arm with a silent mutation to ‘CAG’ to prevent cutting. Insertion locations and orientations were confirmed by PCR (**Figure S1 G S2**).

**Figure 1:**
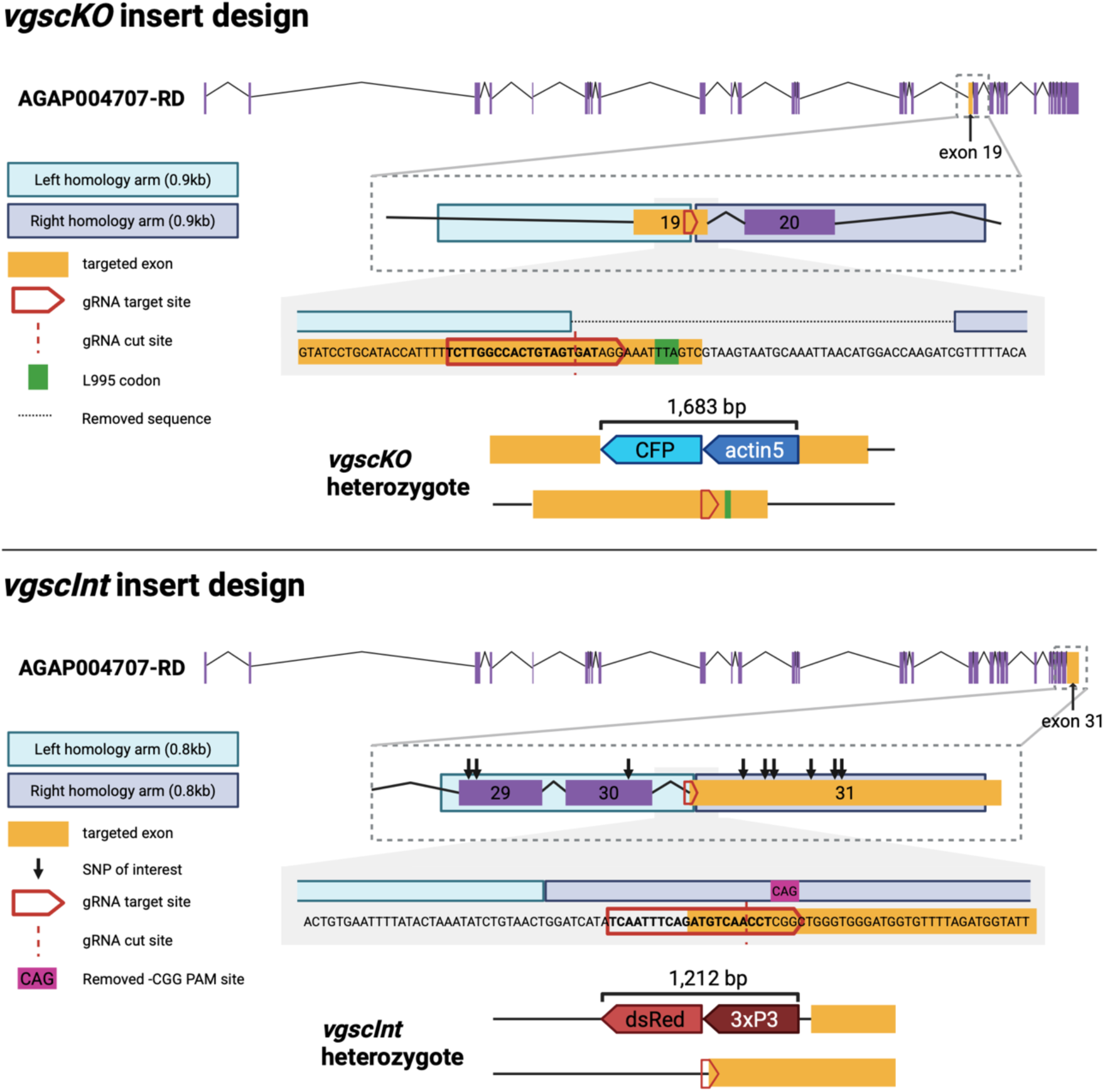
Inserted cassette designs in the *vgscKO* and *vgscInt* lines. The *vgscKO* cassette contains an Actin5c::CFP in reverse orientation relative to *vgsc*, inserted near the 3’ end of exon 19 at 2L:2,422,639 (5’-TCTTGGCCACTGTAGTGAT-3’), close to the important insecticide resistance residue L995, with a 49 bp sequence removed including part of the exon and the beginning of the next intron. The *vgscInt* cassette contains a 3xP3::dsRed in reverse orientation, inserted at the 5’ end of exon 31 at 2L:2,430,607 (5’-(G)TCAATTTCAGATGTCAACCT-3’,) but with a right homology arm recreating the exon to produce an intact *vgsc* coding sequence, with a silent mutation to prevent recutting. Several nearby SNPs of interest as identified in Clarkson *et al. (S)* are indicated, as potential targets for CriMCE editing using *vgscInt* as an intermediate. Diagram created using Biorender.

### Impact of the *vgsc* knockout on fitness and fertility

To assess larval survival, *vgscKO* was backcrossed to G3 to produce known heterozygotes (*vgscKO*-/+); offspring of these heterozygotes were raised to pupation, alongside a G3 control. There was no observed difference in larval survival between the three groups (Binomial GLMM, p > 0.55) (**Figure 2A**).

**Figure 2:**
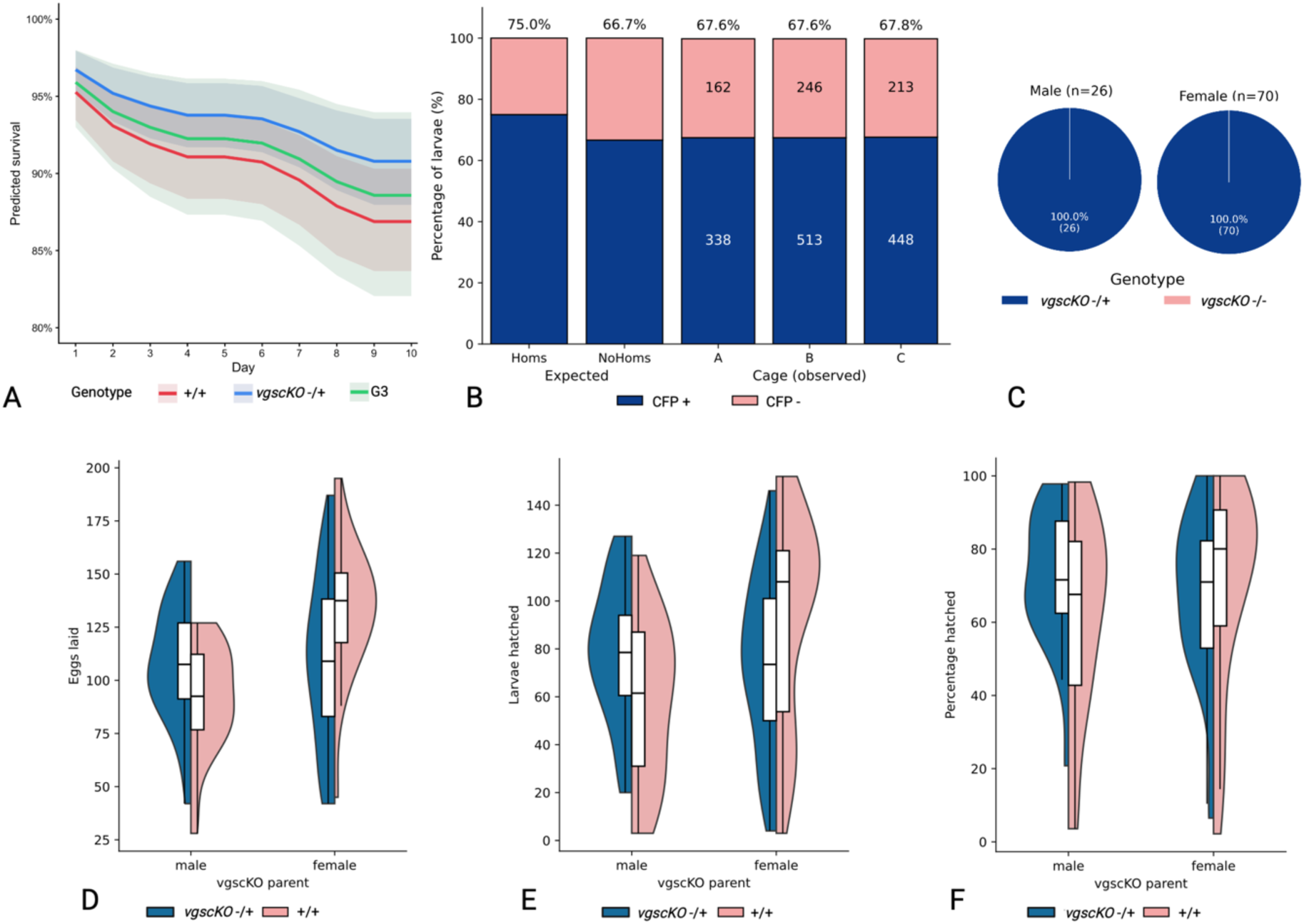
Genotypes and phenotypes of the *vgscKO* colony. A) Hazard predictions for larval survival with three genotypes – *vgscKO-/+*, *+/+* and G3, showing no significant difference in survival between the genotypes. B) Expected and observed percentages of *vgscKO-/+* x *vgscKO-/+* offspring with or without the fluorescent marker; ‘Homs’ indicates the expected ratio of offspring if *vgscKO-/-* homozygotes were viable, and ‘NoHoms’ the expected ratio if they were not viable. C) Observed genotypes of *vgscKO-* parents determined by checking inheritance of the CFP marker in single deposition offspring pools. D), E) and F) Fecundity analyses of *vgscKO-/+* and *+/+* from single depositions of both *vgscKO* females and G3 females crossed to *vgscKO* males.

To determine whether the knockout could be tolerated in homozygosity, *vgscKO*-/+ individuals were crossed together and whole cage clutches were screened for *vgscKO*-proportion by presence of the CFP marker. If *vgscKO*-/- homozygous individuals were viable CFP-positive individuals would be expected to make up 75% of the clutch, whereas if homozygosity was not tolerated the expected CFP-positive proportion would be closer to 67%. In all three repeats CFP-positive individuals made up 67.6-67.8% of the clutch; the observed proportions were 1.75 x 10^11^ more likely to have occurred under the assumption of homozygote non-viability (Binomial likelihood analysis, log_10_ likelihood of 11.24) (**Figure 2B**).

To confirm this lack of homozygotes and to assess impacts of the knockout on fecundity, *vgscKO*-/+ individuals (F_0_) were crossed together and their offspring (F_1_) were blindly backcrossed to G3, with females forced to lay singly to assess clutch phenotype. The *vgscKO* F_1_ were a presumed mixture of knockout homozygous, heterozygous, and wild type; homozygous *vgscKO*-/- individuals would be expected to produce clutches with 100% *vgscKO*- (CFP positive) offspring. All clutches assessed where the knockout was present (n=96) were a mixture of CFP positive and negative, indicating all F_1_ containing the knockout were heterozygous (**Figure 2C**).

The *vgscKO* F1 were assessed for fecundity by counting the number of eggs and larvae produced. Out of 142 single depositions, 13 (11 from *vgscKO* males, two from *vgscKO* females) died before producing eggs and were removed from subsequent analyses. Modelling showed no significant difference between *vgscKO*-/+ and +/+ egg laying (p = 0.58), hatch rate (p = 0.44) or overall larval output (p = 0.82). There was a slight increase in egg batch size from *vgscKO-/+* females compared to *vgscKO-/+* males, but this did not translate into higher larval output.

In summary: no impact of the *vgsc* knockout was observed on fitness or fecundity in heterozygotes, but the knockout does not appear to be tolerated in homozygosity. This indicates that the *vgsc* gene may be haplosufficient in *An. gambiae*.

### Impact of the *vgsc* knockout on insecticide resistance

A blind mixture of *vgscKO*-/+ and +/+ individuals (offspring from *vgscKO*-/+ x *vgscKO*-/+ crosses) were exposed to deltamethrin and DDT, to assess the impact of the knockout on resistance to insecticides which target the VGSC. Post-exposure, *vgscKO* genotype was confirmed by PCR. Permethrin was not considered for use as G3, the base strain of *vgscKO*, shows a low level of resistance to permethrin in our lab colonies.

Exposure of the blinded *vgscKO* to deltamethrin in WHO tubes at the discriminating dose (0.05%) for intervals of 1-15 minutes resulted in 99-100% mortality at all time points (**Figure 3**) demonstrating that even at a reduced dose there is no relevant impact of the knockout on deltamethrin resistance. However, in order to calculate lethal concentrations of both deltamethrin and DDT, WHO bottle bioassays (21) were performed. Individuals from the colony with the knockout (*vgscKO*-/+) were found to have significantly lower overall mortality after low-dose deltamethrin exposure compared to wild-type individuals (*+/+*) (Estimated Marginal Means = −0.92, p = 0.0001), with a resistance ratio of 2.2 at LC50 and 2.9 (**Figure 4**). The response to low-dose DDT was similar (EMM = −0.71, p = <0.0001) but with higher resistance ratios of 3.8 at LC_50_ and 14.4 at LC_99_. Broken down to individual dose level, the lowest dose of deltamethrin showed no difference in mortality between *vgscKO*-/+ and +/+ (p = 0.148), but all subsequent doses showed increasingly large significant differences between the genotypes. Similarly, for DDT the first two doses showed no difference in mortality between genotypes, but all subsequent doses showed significant and increasing differences in mortality (**Table S2**).

**Figure 3:**
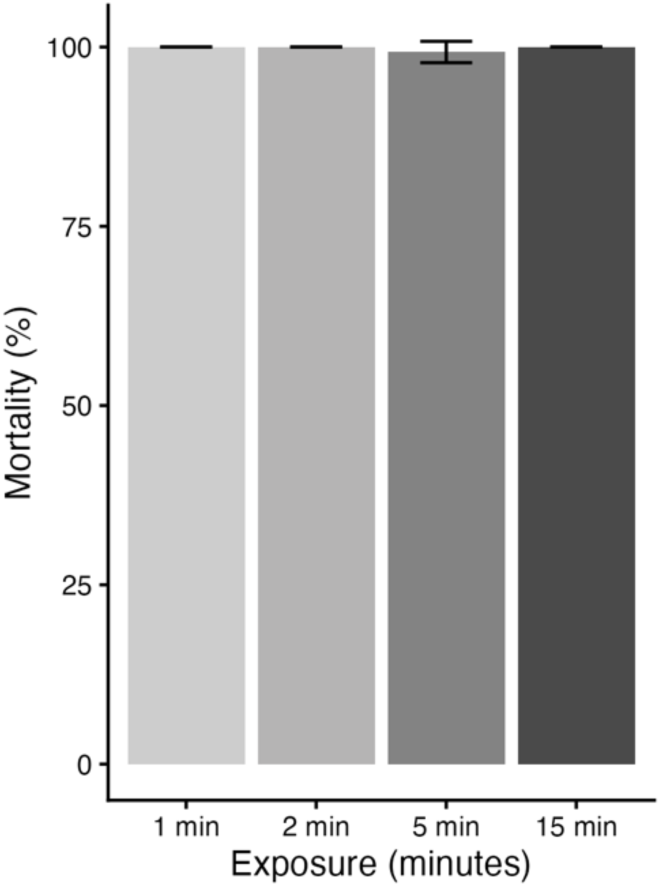
Results from WHO tube test assays where 50/50 mixtures of *vgscKO* -/+ and +/+ females were exposed to the discriminating dose of deltamethrin (0.05%) for four different lengths of time, between 1-15 minutes, with standard deviation error bars.

**Figure 4:**
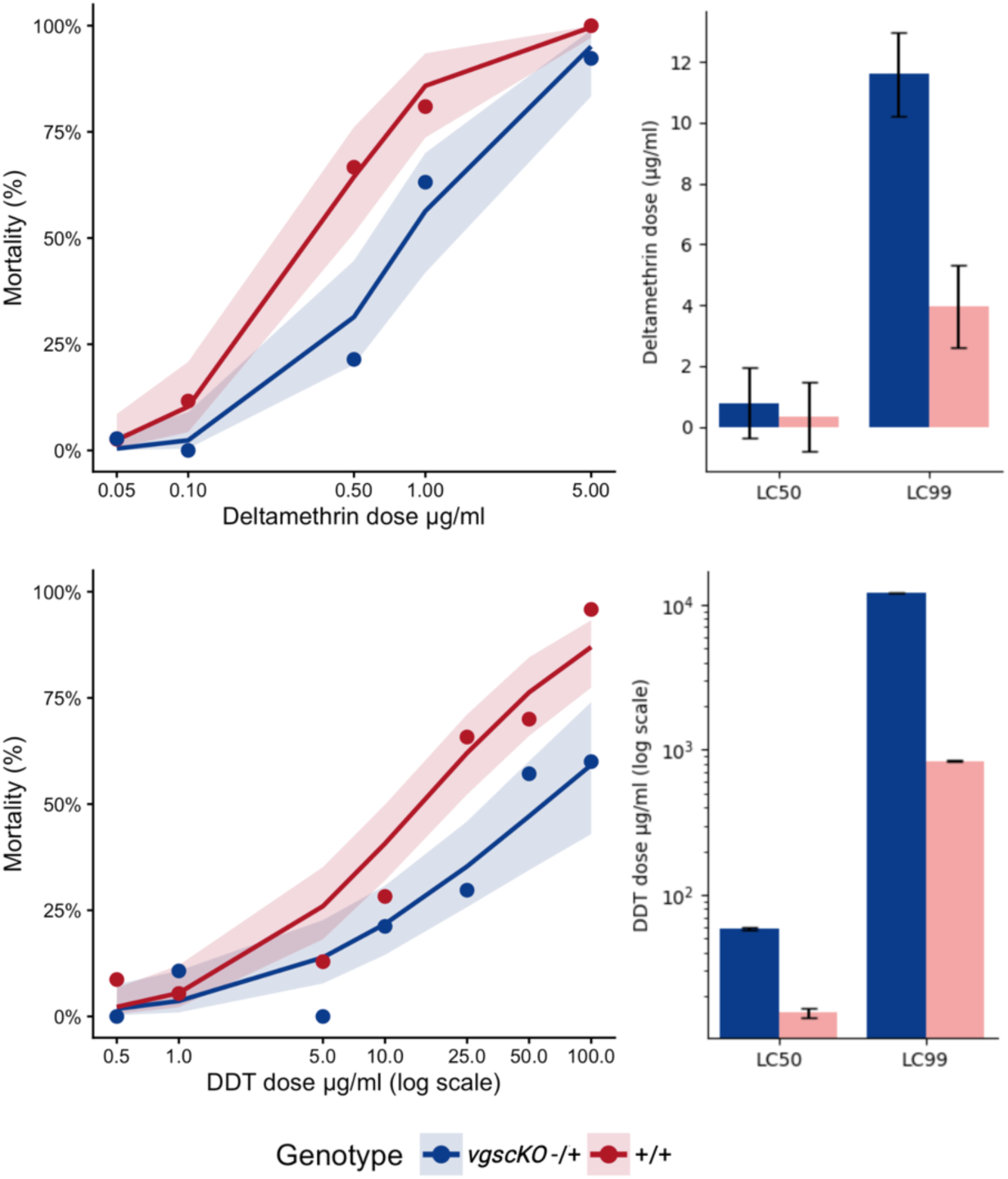
Insecticide response curves and lethal concentrations for deltamethrin and DDT in the *vgscKO* colony. Response curves show true average mortality at each dose and genotype (dots) as well as predicted response curves (lines) with standard error ribbons. LC graphs show LC_50_ and LC_99_ for each genotype and insecticide with standard error bars.

### Intronic marker development for CriMCE in *vgsc*

As the *vgscKO* line demonstrated that *vgsc* knockouts are viable only in heterozygosity, the knockout can be used as a balancer to enrich for or assess the impact of SNPs on the wild type chromosome. In order to produce SNPs via CriMCE, the *vgscInt* line was generated with a different fluorescent marker (RFP) inserted into an intronic region near to where the SNPs of interest occur. In a similar manner to the *vgscKO*, *vgscInt* was not found to be viable in homozygosity despite the insert not impacting any protein coding regions. This was confirmed by crossing *vgscKO* and *vgscInt* together and screening the offspring for both markers; of the 487 larvae screened, none contained both.

## Discussion

### Use of knockouts for investigating *vgsc* function

In this work we have demonstrated that the *vgsc* gene can be knocked out in *Anopheles* with no observable impact on heterozygous fecundity nor larval survival, but that homozygous knockouts are non-viable. This knockout line is therefore ideally suited for use as a balancer allele to evaluate the impact of modifications to *vgsc*. Enhanced genomic surveillance of *Anopheles* has identified multiple SNPs in *vgsc* linked to insecticide resistance; these can be linked through known mechanisms impacting the pyrethroid binding site, through unknown mechanisms elsewhere in the gene, or via linkage disequilibrium with known resistance SNPs (9). With a balancer allele these could be studied in isolation, without the requirement of homozygosity.

The most effective way to insert SNPs as markerless edits is via CriMCE, which ideally requires a fluorescent marker at homozygosity as an intermediate step to be able to observe any successful edit events which result in the desired SNP after a second injection. To achieve this, we designed a fluorescent insert within an intronic region adjacent to an exon with SNPs of interest, hoping to avoid disrupting protein function and therefore produce a homozygous-viable intermediate line. Instead, *vgscInt* was found to not be viable at homozygosity. The *vgsc* gene has a complex set of splice variants and many exons, indicating that the intronic sequences are likely to contain important splice sites and other regulatory regions. While our insert design did not result in sequence removal and kept the coding region intact, increasing the distance between exons or altering the mRNA may have been sufficient to prevent splicing, and we cannot confirm that the insert does not interrupt an essential intronic region. More work is called for to identify splice variants and the factors affecting their production, including better annotation of both introns and exons of the gene.

Despite its inability to exist in homozygosity the *vgscInt* line can be used for CriMCE, albeit in a less efficient manner. After a second injection to remove the fluorescent marker and replace it with the SNP of interest, fluorescence-negative individuals would be screened for the SNP via locked-nucleic acid qPCR in the same way as a straight markerless edit would be. However, by screening out the fluorescent marker half of the unsuccessfully edited chromosomes are removed, enriching the pool for successful edits, and avoiding the ‘needle in a haystack’ search in standard markerless editing. This presents a mechanism for increased editing efficiency in highly conserved genes such as *vgsc*.

### Knockout of *vgsc* and interactions with insecticides

While the *vgscKO* line did not display operationally-relevant insecticide resistance, by reducing the dose of the insecticide it was possible to observe differences in response to both deltamethrin and DDT in heterozygous *vgsc* knockouts. This mirrors previous findings where *vgsc* was knocked down by RNAi in *An. coluzzii* (20). Following RNAi knockdown, resistance to a low dose of deltamethrin increased roughly two-fold in knocked-down individuals compared to controls (20). A similar link between slightly lower *vgsc* transcript level and pyrethroid resistance has also been observed in *Aedes aegypti* (21). We have provided additional evidence that reducing the number of viable copies of *vgsc* decreases the impact of low-level insecticide exposure. Reduced mortality to both DDT and deltamethrin seen in the *vgscKO* line, despite their different VGSC binding profiles, suggests that the knockout reduces the number of VGSCs present on the neurone in a similar manner to an RNAi knockdown.

Despite the variability of bottle assays, which is well documented (22, 23), there was a significant difference detected in the mortality response to both DDT and deltamethrin between the two genotypes, with increased but low-level resistance in *vgscKO*-/+. The cassette inserted into *vgsc* gene likely produces truncated mRNAs which should be degraded by the nonsense-mediated decay pathway (24). We assume that this reduces the level of expression of *vgsc* and therefore the number of VGSCs available for pyrethroid binding, as was seen during RNAi knockdown of *vgsc* previously (20).

Common sense may indicate that a reduced number of insecticide target sites would increase the impact of exposure, as binding sites reach saturation. However, by reducing the dose of insecticide to below saturation point we suggest that a reduced number of VGSCs may actually decrease membrane depolarisation during pyrethroid or DDT exposure, allowing sufficient neuron function to continue transmitting action potentials. While at this point it is not possible to determine the causal mechanism behind fewer VGSC copies reducing pyrethroid mortality, the *vgscKO* line may have use for determining the dynamics of insecticide binding beyond simply being used as a balancer null allele.

In conclusion, we have used gene knockouts to demonstrate that not only is the *vgsc* gene highly conserved, functionally constrained, and of public health relevance through mediating insecticide resistance, but that it is also haplosufficient. This rare combination of features makes *vgsc* simultaneously a mechanistic model system for insecticide resistance and an unusually attractive genetic control target. The two knockout lines produced in this work will facilitate further study of VGSC-pyrethroid dynamics and splice variation, as well as genetic control development.

## Materials and Methods

### Data accessibility

All modelling and figure production code, including package versions, and annotated plasmid sequences have been uploaded to GitHub (25).

### Mosquito strains and rearing

All mosquito strains, and base strains in the case of modified lines, are shown in **Table 2**. Mosquitoes were reared under standard conditions at 27 ± 2°C and 70 ± 10% relative humidity, with a 12:12 light:dark photoperiod which included one-hour simulated dusks and dawns. Larvae were fed with powdered Tetramin® fish food, and adults were fed with 10% sucrose solution *ad libitum*. Blood meals were given before oviposition using donated defibrinated human blood in a Hemotek® membrane feeding system (Hemotek Ltd.). Screening for gene edits via fluorescence was performed at any larval stage for *vgscInt*, as the 3xP3 promoter used in the line is equally visible in the eyes at all stages. For *vgscKO*, larvae were screened approximately six hours after hatching, as this gave the most reliable results with the Actin5c promoter in this line – see **Figure S3** and **Table S3**.

**Table 1.**
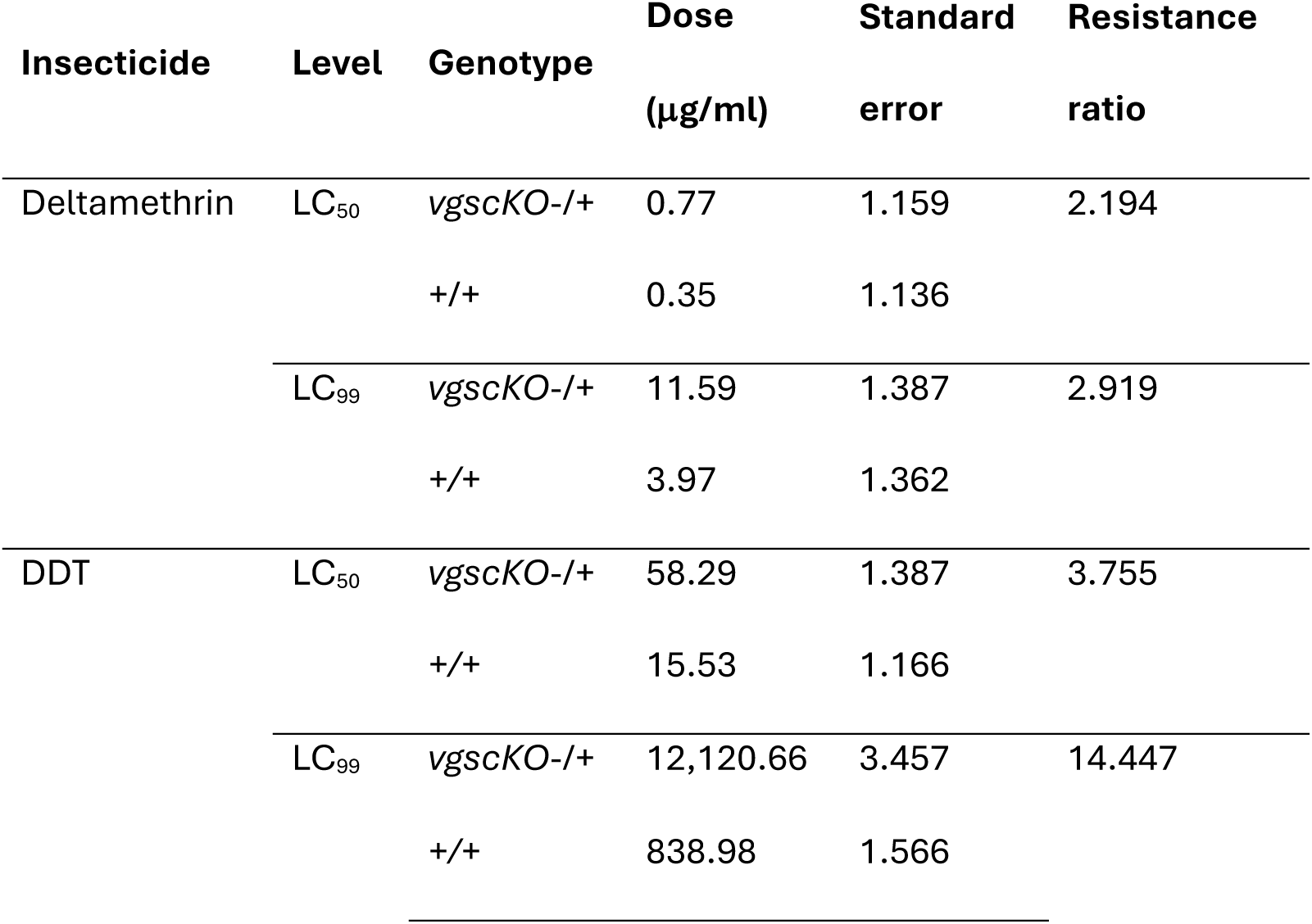
LC_50_ and LC_99_ values and resistance ratios for the *vgscKO* colony, either with (*vgscKO*-/+) or without (+/+) a heterozygous vgsc knockout, after exposure to low doses of deltamethrin or DDT in WHO bottle assays.

**Table 2.** All *Anopheles* strains used or referred to in this work.

| Strain | Species | Modification | Base strain | Ref. |
| --- | --- | --- | --- | --- |
| G3 | *An. gambiae*/*An. coluzzii* hybrid | NA | NA | (14) |
| Kisumu | *An. gambiae* s.s. | NA | NA | Obtained by LSTM from MR4 |
| Kisumu F/F | *An. gambiae* s.s. | Base-change of L995F codon to TTT from TTA, as well as seven synonymous SNPs to prevent self-cutting of the donor plasmid | Kisumu | (17) |
| *vgscKO* | *An. gambiae*/*An. coluzzii* hybrid | Insertion of Actin5C-CFP cassette in exon 19 (plasmids: p20 donor, p21 helper) | G3 | NA |
| *vgscInt* | *An. gambiae* s.s. | Insertion of 3xP3-RFP cassette in intron 30 (plasmids: p111 donor, p110 helper) | Kisumu F/F | NA |

### Microinjections and insert genotyping

Microinjections were performed following the protocol in Fuchs *et al.*(2013) (26); in brief, blood fed females were forced to oviposit and collected eggs were injected in the posterior pole between 30-120 minutes post oviposition. Injection mixes consisted of 200 ng/µL of helper plasmids and 300 ng/µL of donor plasmids in an injection buffer (0.1mM sodium phosphate buffer pH 6.8; 10mM KCl). Injected larvae were screened at the L1 larval stage for transient fluorescence from plasmid expression to confirm injection success and location. Injected larvae were reared and crossed to their respective wild-type strains (**Table 2**), and plasmid integration in F1s was checked by screening for fluorescence either in the eyes (*vgscInt*, 3xP3::RFP) or the midgut (*vgscKO*, Actin5c::CFP).

Insertion location was confirmed by PCR; all DNA extractions were performed by pestle homogenisation of adults in 60 µL and larvae in 40 µL of Lysis Buffer C (200mM Tris pH 8 (Invitrogen, AM9855G), 25 mM EDTA pH 8 (Invitrogen, AM9260G), Tween-20 0.05% (Rockland Inc., TW0020), and Proteinase K 0.4 mg/mL (NEB, P8200G)) (27).

Homogenate was incubated overnight at 56°C and subsequently diluted 1 in 100 (or 1 in 20 for larvae) in molecular-grade water. All PCRs were performed using PrimeSTAR Max DNA polymerase (Takara, R047A) with primers at a final concentration of 0.2 µM, and universal cycling conditions consisting of 30 cycles of: 98°C for 10 seconds, 55°C for 5 seconds, and 72°C for 90 seconds, with a final indefinite hold at 12°C (**Figure S1** and **Table S4**). The same protocol was used for all subsequent genotyping.

### Larval survival assay

To determine the hazard ratio associated with the *vgscKO* knockout during larval development, *vgscKO* L1 stage larvae were screened for the insert on the first day after hatching and positives and negatives were separated into eight trays, with four smaller trays (8 x 10 cm) containing 25 larvae and four larger trays (12 x 15 cm) containing 50 larvae for each genotype. Two trays of G3, one small tray of 25 larvae and one large tray of 50, were taken at the same age and on the same day to act as a control. The number of remaining larvae in each tray was counted each day for ten days.

Daily mortality risk of the three genotypes (*vgscKO*-/+, +/+, and G3) was determined using a binomial generalised linear mixed model with a complementary log-log (cloglog) link, allowing regression coefficients to be interpreted as log-hazard ratios. This approach was chosen over a Cox proportional hazards approach as the Genotype, Tray size (either 25 larvae or 50 larvae), and Day were included as fixed effects; individual tray identity was included as a random intercept. Models were fitted by maximum likelihood using the *lme4* package in R (version 4.5.2). Post-hoc testing was performed with the *DHARMa* package to validate the model, with a final simplified model chosen removing non-significant factors.

Daily hazard estimates were produced by applying the inverse complementary log-log transformation to the linear predictor of the model, with hazards averaged across tray sizes as these were not a significant factor in the model, and the cumulative product of daily survival probabilities was calculated for each day (1–10). Parametric bootstrapping was performed with 1,000 replicates using the *bootMer* function in *lme4*, producing hazard and survival curves for all genotype x day combinations, and pointwise 95% confidence intervals. Curves were plotted with confidence interval bands using *ggplot2*. Raw data can be found in **Table S5**, and scripts for models and plots on GitHub (25).

### Single depositions for fecundity and screening for homozygotes

To produce a mixed batch of *vgscKO* heterozygotes, homozygotes and wild-type offspring, *vgscKO* larvae were purified for the insert and raised to pupation. Individual pupae were allowed to emerge singly in cups, and we removed legs through the cup mesh for individual genotyping by PCR to confirm the results of the fluorescent screening. Positive individuals (approximately 20 males and 20 females) were mixed together and allowed tomate for one week before blood-feeding; F1 offspring (a blind mix of heterozygotes, homozygotes, and wild type) were sexed at the pupal stage and split into two cages to mate with opposite sex G3. Females from each cage were separated to oviposit in single deposition: a total of 70 G3 females mated to unknown genotype *vgscKO* males, and 69 *vgscKO* females mated to G3 males. Eggs and hatched larvae were counted, and offspring pools were checked for the fluorescent marker. A mixture of positives and negatives in a single pool indicated a heterozygous *vgscKO* parent, a pool of 100% positives indicated a homozygous *vgscKO* parent, and a pool entirely absent of the *vgscKO* marker indicated a wild-type parent. Any females which did not produce eggs were checked for mating status by spermathecae dissection and microscopy (**Table S6**). Without counting eggs or larvae, we screened an additional 36 *vgscKO* females for homozygosity which were positive for the insert but with unknown zygosity. (**Table S7**).

The impact of genotype on egg production, larval output and hatch rate was modelled using either negative binomial or quasibinomial generalised linear modelling, with models tested for fit using post-hoc analyses. Estimated marginal means were calculated using the *emmeans* package. Violin plots and pie charts were produced in Python (version 3.14.2) using matplotlib. Full model details and plot code are hosted on GitHub (25).

### Group oviposition to screen for homozygotes

Mixed batches of *vgscKO* heterozygotes, homozygotes and wild-type offspring were produced as above and screened at L1 stage to determine the proportion of the fluorescent marker in each egg batch. This was repeated with three separate generations, with each generation blood fed two to three times to collect multiple egg papers, to maximise the number of larvae screened for each cage. A total of 500, 759, and 661 larvae were screened from each generation (**Table S8**).

To determine the likelihood of viable homozygotes in the observed proportions of CFP-positive *vgscKO*-/+ x *vgscKO*-/+ offspring, we performed a binomial likelihood analysis. If homozygotes were viable, 75.0% of the offspring in each pool would contain the marker, and if they were not viable 66.7% would contain the marker. The log-likelihoods for both hypotheses were generated, and a likelihood ratio was calculated. Full model details and plot code are hosted on GitHub (25).

### *vgscKO* x *vgscInt* crosses to screen for homozygotes

Positive individuals from both colonies (screened by fluorescence) were sexed as pupae and crossed together, with one cage of *vgscKO* females x *vgscInt* males, and one cage with *vgscKO* males x *vgscInt* females, with approximately 20 of each genotype per cage. Cages were blood fed twice and allowed to oviposit en masse twice; larvae were screened on day 1 post-hatching for both fluorescent markers.

### Insecticide assays

Known *vgscKO-/*+ adults were mated en masse to produce mixed offspring; offspring were screened for the *vgscKO* insert and raised in trays of 65 *vgscKO-/*+ and 65 *+/+* to produce cages of 50/50 positives and negatives, for blind testing on insecticides. We transferred mosquitoes to paper cups with 25 females each and moved to the insectary testing suite to acclimatise for one hour, before exposure. Mosquitoes were allowed to feed on sucrose solution ad libitum before and after exposure. For all insecticide testing, knockdown was scored at one hour post exposure and mortality was scored at 24-hours post exposure. At the 24-hour point, mosquitoes from each replicate were separated by dead or alive status and retained in individual wells of 96-well plates for genotyping.

Blind mixes were first tested on WHO paper containing the discriminating dose of deltamethrin (0.05%) for 15 minutes, otherwise following standard WHO tube testing procedure (28). Mosquitoes were exposed to six tubes containing deltamethrin papers (WHO, batch DE 1165) and three containing control papers (WHO, batch PY 475). As mortality was 100% in all six test tubes, no genotyping was performed (**Table S1**).

Blinded *vgscKO* mixes were tested against deltamethrin and DDT in bottle assays, following WHO bottle assay standard procedures (22), with exposure times of one hour for both insecticides. Dilutions of deltamethrin (Sigma Aldrich, 45423) and DDT (Sigma Aldrich, 31041) were produced from powdered stocks in acetone (Sigma Aldrich, 179124); 1 ml of each concentration was used to coat 250 mL Wheaton glass bottles by hand turning. Deltamethrin bottles were wrapped in aluminium foil and stored at 4°C for up to 7 days before use. DDT bottles were coated and used the same day to avoid insecticide breakdown. Insects were exposed and genotyped as described above (**Table SG**).

Insecticide effect was analysed at all doses using a generalised linear model with a probit link, including individual bottles as a random effect, using the *glmer* package. Model fit was checked and a pairwise emmeans comparison was performed using the *emmeans* package to produce a contrast between genotypes for the overall insecticide mortality as well as mortality at each individual dose (**Tables S2**). A prediction grid, with standard errors, was produced using the *lme4* package, and used to plot predicted mortality, with standard error ribbons, using *ggplot2*. Finally, lethal concentration tables were produced for each genotype of each insecticide using the *ecotox* package, and displayed as bar charts with standard error bars produced using *matplotlib*. Full model details and plot code are hosted on GitHub (25).

## Supplementary Material

**Figure S1:**
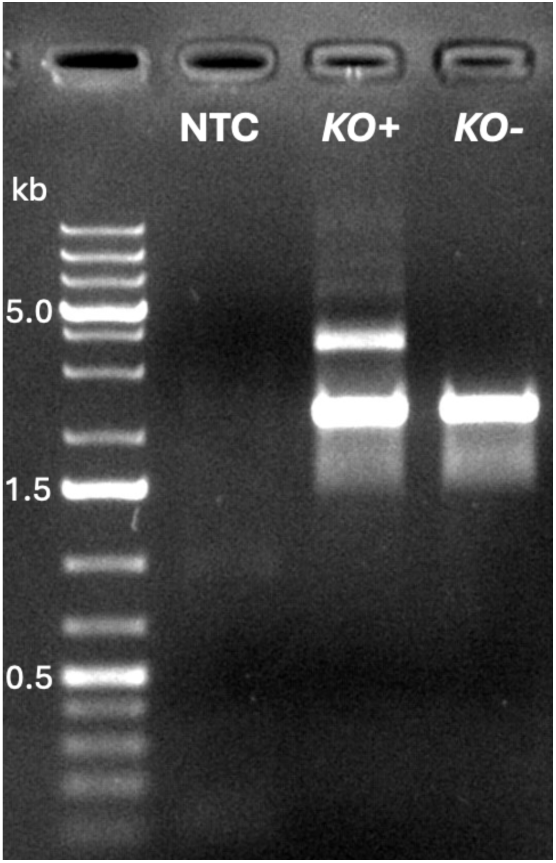
PCR to confirm insertion of the *vgscKO* cassette at the target site. Primers used: TN149 and TN48, which bind on the genomic DNA outside of the plasmid homology arms. Expected band sizes are 3,899 bp with the *vgscKO* cassette, 2,264 bp for the wild type. DNA ladder is GeneRuler 1kb plus (ThermoFisher). NTC: non-template control. KO+: *vgscKO*-/+ (with the insert). KO-: *+/+* (wild type).

**Figure S2:**
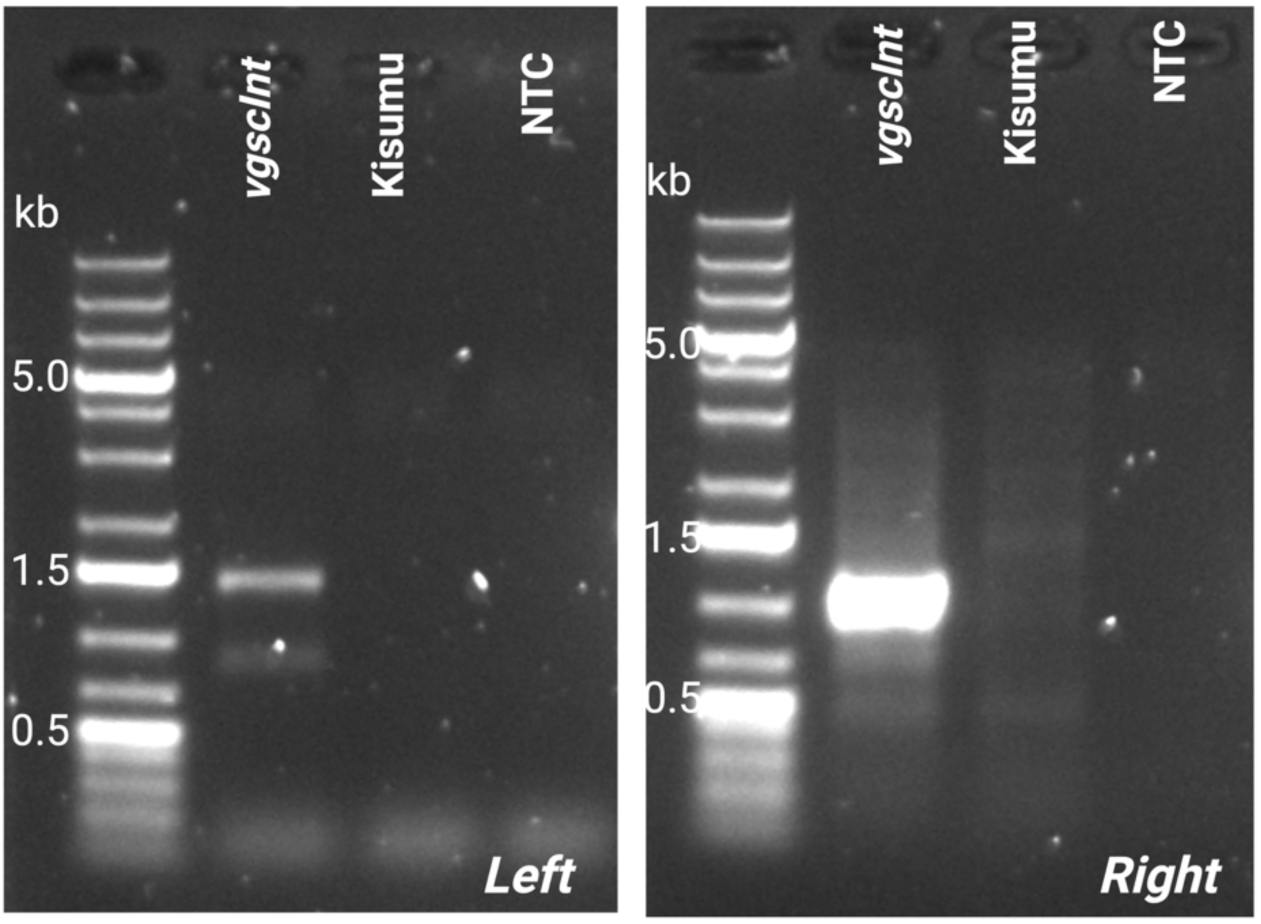
PCRs to confirm insertion of the *vgscInt* cassette at the target site, with Kisumu DNA as an extra control. Primers used for the left side of the insert: CriMCE PH PresAbs1 fwd and TN8, which bind on the genomic DNA outside of the left homology arm and on the SV40 terminator within the cassette, with an expected band size of 1,482 bp. Primers used for the right side of the insert: TN540 and TN957, which bind on the 3xP3 promoter within the cassette and on the genomic DNA outside of the right homology arm, with an expected band size of 1,081 bp. DNA ladder is GeneRuler 1kb plus (ThermoFisher). NTC: non-template control.

**Figure S3:**
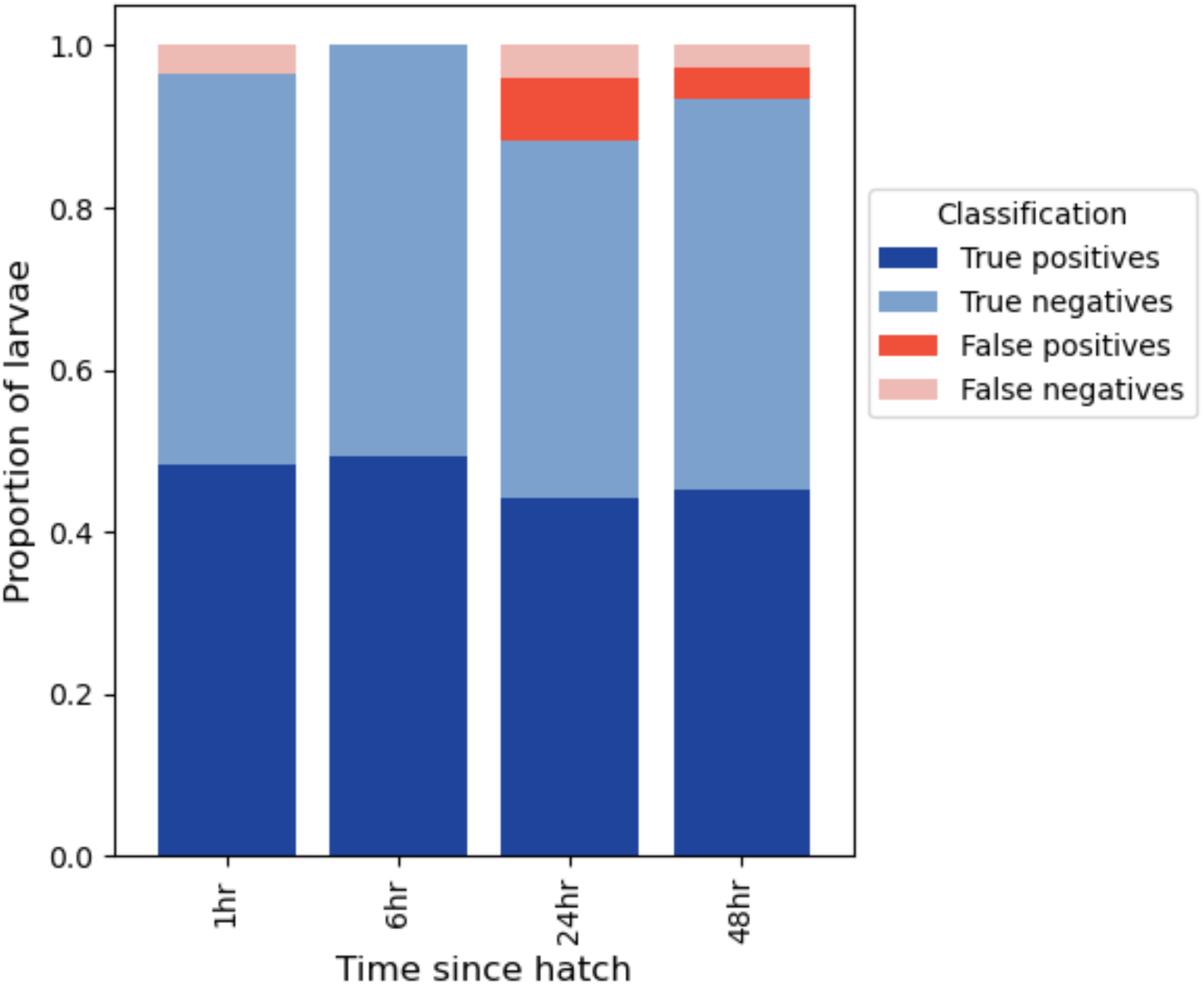
Accuracy of *vgscKO* fluorescent screening at different ages post hatching. At each time point, 94 individuals were screened first for presence of the CFP marker and subsequently genotyped by PCR to determine true genotype.

---

**Table S1:** WHO tube test raw data of blind mixed populations of *vgscKO (vgscKO*-/+ and +/+), with six tubes exposed to Deltamethrin 0.05% papers for 15 minutes, and two tubes exposed to PY Control papers for 15 minutes. Deltamethrin-exposed mosquitoes had 100% mortality rates in all tubes.

**Table S2:** Estimated marginal means for response to different doses of deltamethrin and DDT in WHO bottle assays, showing probit estimates of response differences, standard errors and p values at each dose.

**Table S3:** Raw data from optimum screening age experiment, showing the total number of larvae at each age found to contain the *vgscKO* marker (CFP+) or not (CFP-) by fluorescent microscopy, and their subsequent true genotype determined by PCR. Any PCR result which was potentially ambiguous was discarded as ‘inconclusive’ to ensure accuracy of the PCR assay in determining genotype.

**Table S4:** Primer sequences for confirming insert and determining genotype of *vgscInt* and *vgscKO*, with band sizes given for insert positive and negative samples (note: heterozygotes have both bands). All primers bind outside of the homology arms of the donor plasmid used to create the lines, to ensure correct situation of the donor sequence.

**Table S5:** Raw data from larval survival assay, with the number of larvae alive in each tray and percent survival from the original number of larvae on Day 1, as well as the life stage of the majority of larvae in each tray.

**Table S6:** Raw data from the single female depositions of *vgscKO* x G3 mating pairs, showing the number of eggs and larvae per female and the inferred genotype of the *vgscKO* parent based on the proportion of offspring expressing the fluorescent marker. Females which died before laying, if possible, were dissected to examine their spermathecae for mating status; ‘pos’ indicates sperm were present in the spermathecae, ‘neg’ indicates sperm were absent.

**Table S7:** Raw data from single depositions where the parent genotype was determined by the proportion of offspring containing the fluorescent marker; some of these are counted from the single deposition data in Table S6. Any larval pools with fewer than 10 living larvae were discarded.

**Table S8:** Raw data from three cage depositions of *vgscKO*-/+ x *vgscKO*-/+ crosses where the pooled offspring were screened for presence of the fluorescent marker.

**Table S9:** Raw data from WHO bottle assays where blinded mixed cohorts of *vgscKO* were exposed to deltamethrin and DDT.

## References

1. World Health Organization. World malaria report 2025: Addressing the threat of antimalarial drug resistance. Geneva; 2025.

2. Pryce J, Richardson M, Lengeler C. Insecticide-treated nets for preventing malaria. Cochrane Database Syst Rev. 2018;11(11):Cd000363.

3. Zaim M, Aitio A, Nakashima N. Safety of pyrethroid-treated mosquito nets. Medical and Veterinary Entomology. 2000;14(1):1–5.

4. Davies TG, Field LM, Usherwood PN, Williamson MS. DDT, pyrethrins, pyrethroids and insect sodium channels. IUBMB Life. 2007;59(3):151–62.

5. Davies TG, Field LM, Usherwood PN, Williamson MS. A comparative study of voltage-gated sodium channels in the Insecta: implications for pyrethroid resistance in Anopheline and other Neopteran species. Insect Mol Biol. 2007;16(3):361–75.

6. Dong K, Du Y, Rinkevich F, Nomura Y, Xu P, Wang L, et al. Molecular biology of insect sodium channels and pyrethroid resistance. Insect Biochem Mol Biol. 2014;50:1–17.

7. Yu FH, Catterall WA. Overview of the voltage-gated sodium channel family. Genome Biology. 2003;4(3):207.

8. Olson ROD, Liu Z, Nomura Y, Song W, Dong K. Molecular and functional characterization of voltage-gated sodium channel variants from *Drosophila melanogaster*. Insect Biochemistry and Molecular Biology. 2008;38(5):604–10.

9. Clarkson CS, Miles A, Harding NJ, O’Reilly AO, Weetman D, Kwiatkowski D, et al. The genetic architecture of target-site resistance to pyrethroid insecticides in the African malaria vectors *Anopheles gambiae* and *Anopheles coluzzii*. Molecular Ecology. 2021;30(21):5303–17.

10. Davies TE, O’Reilly AO, Field LM, Wallace B, Williamson MS. Knockdown resistance to DDT and pyrethroids: from target-site mutations to molecular modelling. Pest Manag Sci. 2008;64(11):1126–30.

11. Miles A, Harding NJ, Bottà G, Clarkson CS, Antão T, Kozak K, et al. Genetic diversity of the African malaria vector *Anopheles gambiae*. Nature. 2017;552(7683):96–100.

12. Burt A. Site-specific selfish genes as tools for the control and genetic engineering of natural populations. Proceedings of the Royal Society of London Series B: Biological Sciences. 2003;270(1518):921–8.

13. Kyrou K, Hammond AM, Galizi R, Kranjc N, Burt A, Beaghton AK, et al. A CRISPR– Cas9 gene drive targeting doublesex causes complete population suppression in caged *Anopheles gambiae* mosquitoes. Nature Biotechnology. 2018;36(11):1062–6.

14. Hammond A, Galizi R, Kyrou K, Simoni A, Siniscalchi C, Katsanos D, et al. A CRISPR-Cas9 gene drive system targeting female reproduction in the malaria mosquito vector *Anopheles gambiae*. Nature Biotechnology. 2016;34(1):78–83.

15. Tapia A, Giachello CN, Palomino-Schätzlein M, Baines RA, Galindo MI. Generation and Characterization of the *Drosophila melanogaster paralytic* Gene Knock-Out as a Model for Dravet Syndrome. Life. 2021;11(11):1261.

16. Bosch JA, Birchak G, Perrimon N. Precise genome engineering in *Drosophila* using prime editing. Proceedings of the National Academy of Sciences. 2021;118(1):e2021996118.

17. Grigoraki L, Cowlishaw R, Nolan T, Donnelly M, Lycett G, Ranson H. CRISPR/Cas9 modified *An. gambiae* carrying kdr mutation L1014F functionally validate its contribution in insecticide resistance and combined effect with metabolic enzymes. PLOS Genetics. 2021;17(7):e1009556.

18. Kistler KE, Vosshall LB, Matthews BJ. Genome engineering with CRISPR-Cas9 in the mosquito *Aedes aegypti*. Cell reports. 2015;11(1):51–60.

19. Morianou I, Crisanti A, Nolan T, Hammond AM. CRISPR-Mediated Cassette Exchange (CriMCE): A Method to Introduce and Isolate Precise Marker-Less Edits. The CRISPR Journal. 2022;5(6):868–76.

20. Nardini L, Brito-Fravallo E, Campagne P, Pain A, Genève C, Vernick KD, et al. The voltage-gated sodium channel, *para*, limits *Anopheles coluzzii* vector competence in a microbiota dependent manner. Scientific Reports. 2023;13(1):14572.

21. Kubik TD, Snell TK, Saavedra-Rodriguez K, Wilusz J, Anderson JR, Lozano-Fuentes S, et al. *Aedes aegypti* miRNA-33 modulates permethrin induced toxicity by regulating VGSC transcripts. Scientific Reports. 2021;11(1):7301.

22. Corbel V, Kont MD, Ahumada ML, Andréo L, Bayili B, Bayili K, et al. A new WHO bottle bioassay method to assess the susceptibility of mosquito vectors to public health insecticides: results from a WHO-coordinated multi-centre study. Parasites & Vectors. 2023;16(1):21.

23. Althoff RA, Huijben S. Comparison of the variability in mortality data generated by CDC bottle bioassay, WHO tube test, and topical application bioassay using Aedes aegypti mosquitoes. Parasites & Vectors. 2022;15(1):476.

24. Gatfield D, Unterholzner L, Ciccarelli FD, Bork P, Izaurralde E. Nonsense-mediated mRNA decay in *Drosophila*: at the intersection of the yeast and mammalian pathways. Embo j. 2003;22(15):3960–70.

25. Pescod P. vgscKOInt-Scripts - https://github.com/poppy541/vgscKOInt-Scripts. GitHub; 2026.

26. Fuchs S, Nolan T, Crisanti A. Mosquito transgenic technologies to reduce *Plasmodium* transmission. Methods Mol Biol. 2013;923:601–22.

27. Korlević P, McAlister E, Mayho M, Makunin A, Flicek P, Lawniczak MKN. A Minimally Morphologically Destructive Approach for DNA Retrieval and Whole-Genome Shotgun Sequencing of Pinned Historic Dipteran Vector Species. Genome Biology and Evolution. 2021;13(10):evab226.

28. World Health Organization. Standard operating procedure for testing insecticide susceptibility of adult mosquitoes in WHO tube tests. Geneva; 2022.

